# SARS-CoV-2 infects and replicates in photoreceptor and retinal ganglion cells of human retinal organoids

**DOI:** 10.1101/2021.10.09.463766

**Authors:** Yotam Menuchin-Lasowski, André Schreiber, Aarón Lecanda, Angeles Mecate-Zambrano, Linda Brunotte, Olympia E. Psathaki, Stephan Ludwig, Thomas Rauen, Hans R. Schöler

## Abstract

Several studies have pointed to retinal involvement in COVID-19 disease, yet many questions remain regarding the ability of SARS-CoV-2 to infect and replicate in retinal cells and its effects on the retina. Here we have used human stem cell–derived retinal organoids to study retinal infection by the SARS-CoV-2 virus. Indeed, SARS-CoV-2 can infect and replicate in retinal organoids, as it is shown to infect different retinal lineages, such as retinal ganglion cells and photoreceptors. SARS-CoV-2 infection of retinal organoids also induces the expression of several inflammatory genes, such as interleukin 33, a gene associated with acute COVID-19 disease and retinal degeneration. Finally, we show that the use of antibodies to block the ACE2 receptor significantly reduces SARS-CoV-2 infection of retinal organoids, indicating that SARS-CoV-2 infects retinal cells in an ACE2-dependent manner. These results suggest a retinal involvement in COVID-19 and emphasize the need to monitor retinal pathologies as potential sequelae of “long COVID”.

## Introduction

The coronavirus disease 2019 (COVID-19), caused by the severe acute respiratory syndrome coronavirus 2 (SARS-CoV-2), is a global pandemic that is responsible for millions of fatalities. While mild-to-severe respiratory symptoms are most commonly associated with the disease (Huang et al., 2020), neurological (Asadi-Pooya and Simani, 2020; Mao et al., 2020) and ocular (Wu et al., 2020) symptoms have also been described in patients.

Growing evidence suggests retinal involvement in some cases of COVID-19. Different reports identified SARS-CoV-2 RNA in retinal and optic nerve biopsies taken from deceased COVID-19 patients (Casagrande et al., 2020; Casagrande et al., 2021), while different retinal anomalies were also identified in patients (Burgos-Blasco et al., 2020; Conrady et al., 2021; Marinho et al., 2020; Pereira et al., 2020; Rodriguez-Rodriguez et al., 2021; Virgo and Mohamed, 2020). Yet, retinal involvement in COVID-19 remains a controversial topic, as other studies were not able to detect retinal pathologies in patients (Pirraglia et al., 2020). Furthermore, one study reported failure to isolate virus from SARS-CoV-2 RNA–positive retinal biopsies and an inability to detect any SARS-CoV-2 spike protein from those biopsies by immunohistochemical analysis, leading investigators to suggest that SARS-CoV-2 can infect but not actively replicate in retinal cells (Casagrande et al., 2021). It also remains unclear which retinal structures are infected by SARS-CoV-2 and whether the retinal anomalies reported are due to retinal infection or systemic organ dysfunction (Casagrande et al., 2020; de Figueiredo et al., 2020).

Organoids are three-dimensional, tissue cultures that resemble specific organs in their morphology, cell-type composition, or function. Different research groups have used organoids generated from human stem cells to study the infection of SARS-CoV-2 in the context of different organs (Lamers et al., 2020; Monteil et al., 2020; Pellegrini et al., 2020; Ramani et al., 2020; Zhang et al., 2020). Such studies have demonstrated that SARS-CoV-2 can infect neurons and neural progenitors (Ramani et al., 2020; Zhang et al., 2020). Retinal organoids are among the most physiologically accurate neural organoid models, as they contain all the major retinal cell types, follow a developmental trajectory similar to that of the human retina *in vivo*, and are correctly organized in a typical layered structure (Achberger et al., 2019; Nakano et al., 2012; Zhong et al., 2014).

Here we use retinal organoids to show that SARS-CoV-2 can infect retinal cells, mainly retinal ganglion cells (RGCs) but also photoreceptors, as well as replicate in them. Moreover, retinal SARS-CoV-2 infection induces the expression of several inflammatory genes and is reduced by treatment of the organoids with an anti-ACE-2 antibody.

## Results

### SARS-CoV-2 infects retinal organoids and replicates in retinal cells

To generate retinal organoids in large quantities we have combined different widely used protocols (Capowski et al., 2019; Kuwahara et al., 2015; Zhong et al., 2014). The organoids generated followed correct retinal development and contained a thick layer of VSX2+ retinal progenitors and a basal layer of differentiated PAX6+/VSX2-RGC and amacrine cells by day 50 into the protocol (Figure 1A). At later stages, the organoids had gained a layered retinal structure with an inner nuclear layer (INL), containing the different INL cell types such as OTX2+ bipolar cells, AP2A+ amacrine and horizontal cells, and CRALBP+ Müller glia, separated from an outer nuclear layer (ONL), containing OTX2+ photoreceptors (Figure 1B). The photoreceptor cells expressed rod and cone opsins concentrated in outer segment–like structures (Figure 1C), indicating highly mature organoids.

**Figure 1:**
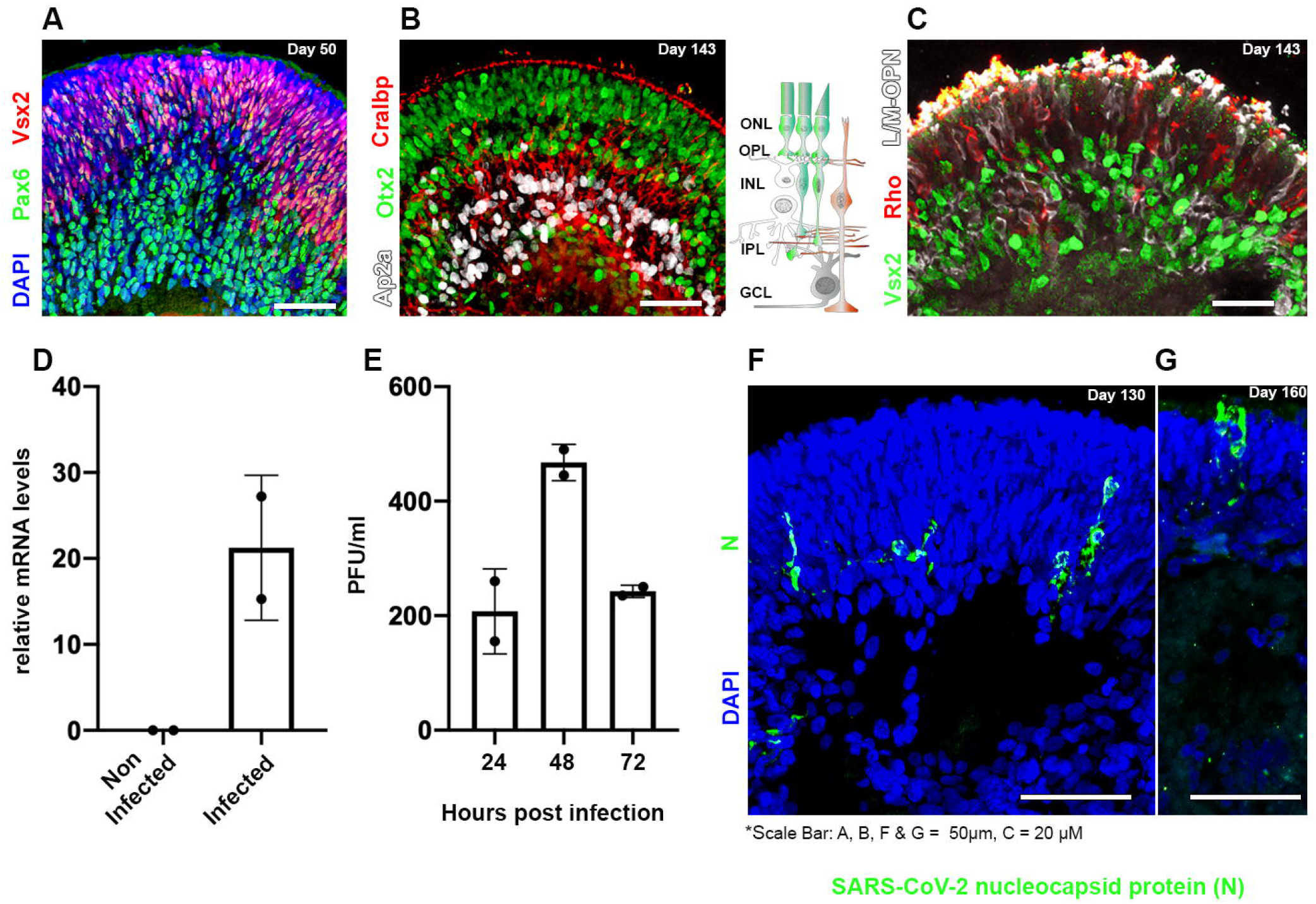
Human iPSC-derived retinal organoids can be infected by SARS-CoV-2. Retinal organoids on day 50 of differentiation are shown to contain VSX2+ (green) retinal progenitors and PAX6+ (white)/VSX2-amacrine and retinal ganglion cells (RGCs) **(A)**. On day 143 of differentiation, the organoids are shown to be organized in a layered structure and to contain AP2a+ (white) amacrine and horizontal cells, CRALBP+ (red) Müller glia cells, and OTX2+ (green) cells **(B).** Th**e** sketch in (**B**) depicts the cell types and structure of the vertebrate retina. ONL: outer nuclear layer; OPL: outer plexiform layer; INL: inner nuclear layer; IPL: inner plexiform layer; GCL: ganglion cell layer. On day 143 of differentiation, the photoreceptors in the organoids are stained against the photosensitive proteins L/M-OPN (white) and RHODOPSIN (RHO, red) **(C)**. Real-time PCR identified SARS-CoV-2 genomic RNA within retinal organoids treated with SARS-CoV-2 but not in controls on day 125 **(D)**. A viral plaque assay was used to assess viral titrations in SARS-CoV-2 treated organoids on day 125 **(E).** Each repeat (N=2) in **(D)** and **(E)**, is a separate group of five organoids infected or not infected in a separate well. Immunofluorescence analysis was used to detect SARS-CoV-2 nucleocapsid (N)-positive cells in infected organoids at day 130 **(F)** or 160 **(G)** of differentiation.

To test the ability of SARS-CoV-2 to infect retinal cells, relatively mature human retinal organoids (day of differentiation ≥ 125) were treated with SARS-CoV-2 and then incubated for different incubation periods. The infected organoids were then analyzed by different methods to assess SARS-CoV-2 infection and replication. SARS-CoV-2 RNA was detected by qPCR analysis in SARS-CoV-2–treated organoids but not in the controls (Figure 1D), suggesting that the treated organoids were indeed infected with SARS-CoV-2. Furthermore, a viral plaque assay was used to measure the active viral concentrations produced by the infected organoids after different incubation times. The appearance of viral plaques in the cell cultures treated with supernatant that was incubated for 24 hours with the infected organoids indicated that new virus progeny was being generated, providing the first evidence that SARS-CoV-2 replicates within human retinal tissue. Accordingly, virus titers were increased after 48 hours of incubation with the infected organoids but surprisingly were reduced back to levels which are similar to the 24-hour time point after 72 hours. A similar phenomenon was reported in SARS-CoV-2–infected brain organoids (Zhang et al., 2020).

Finally, an immunofluorescence analysis identified a small number of SARS-CoV-2 nucleocapsid protein (N) positive cells in SARS-CoV-2–treated organoids on days 130 and 160 of differentiation (Figure 1F and G) but not in controls (Figure 2F). Interestingly, many of the N+ cells were found in the ONL of the organoid and some exhibited a typical photoreceptor morphology (Figure 1G). While most of the work presented here was made using human induced pluripotent stem cells (hiPSCs), organoids derived from human embryonic stem cells were also found to be permissive to infection (Figure S1A).

**Figure 2:**
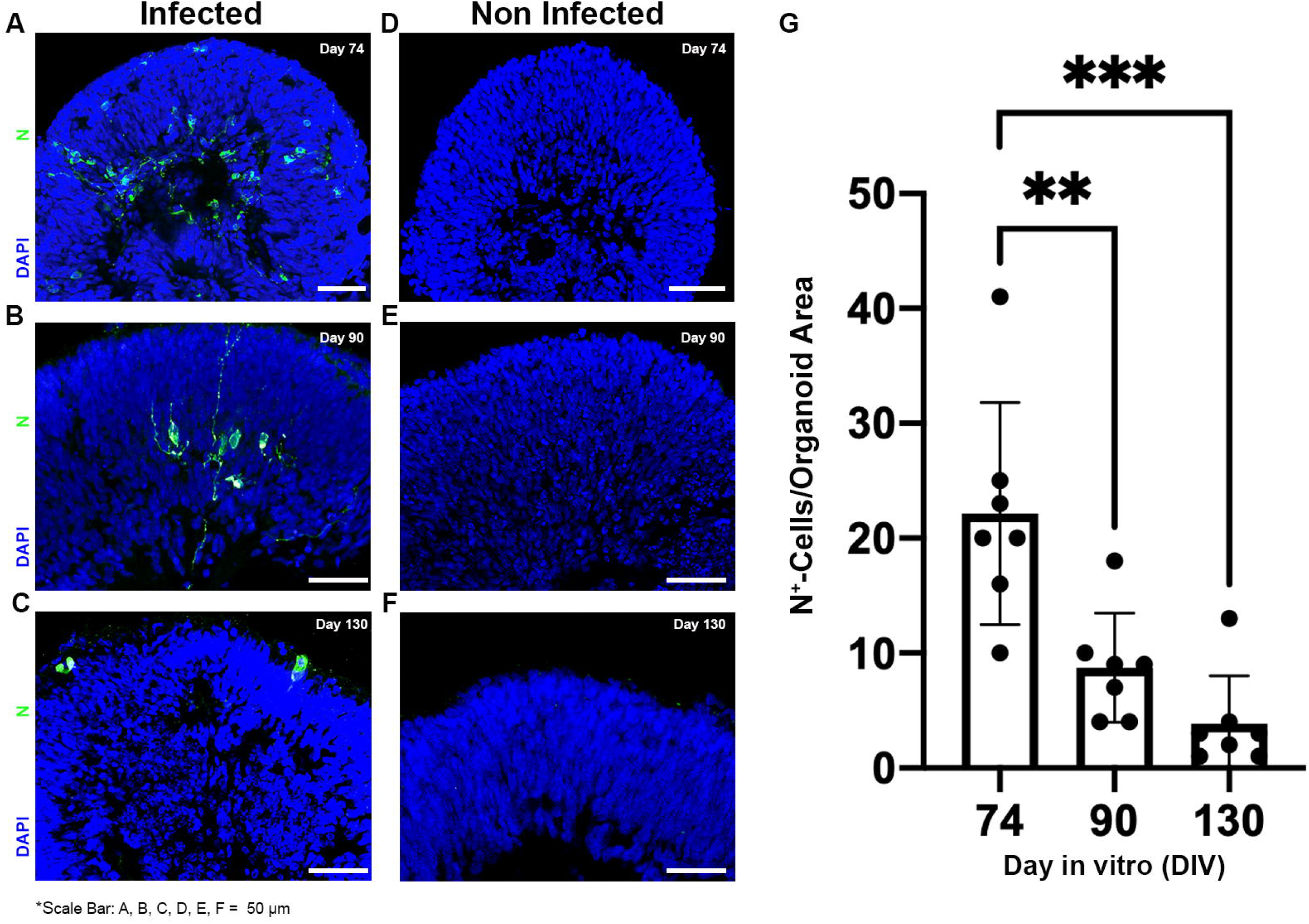
SARS-CoV-2 nucleocapsid-positive cells are more abundant in retinal organoids infected at an earlier stage of differentiation. SARS-CoV-2 nucleocapsid (N) immunostaining was used to quantify the number of N+ (green) cells in retinal organoids that were infected at three different time points: day 74 **(A)**, day 90 **(B)**, and day 130 **(C)**. An age-matched noninfected control for each stage: day 74 **(D),** Day 90 **(E)**, and day 130 **(F)** is also presented. All organoids were incubated for 96 hours after infection and before fixation. The number of N+ cells counted in the organoids of each stage was compared **(G)**. N=7 organoids, taken from two different infected or control wells, ANOVA P-value = 0.0002, F = 14.15; **: P-value = 0.0038; ***: P-value = 0.0002.

The number of N+ cells identified in retinal organoids at these late developmental stages was relatively low (3.8 per organoid, on average, on day 130, Figure 2D). To investigate whether retinal organoids at earlier developmental stages are more susceptible to SARS-CoV-2 infection, we infected less mature retinal organoids, at either day 74 or day 90, and compared the number of N+ cells with SARS-CoV-2–infected 130-day-old retinal organoids (Figure 2A, B and C). The number of N+ cells on day 74–infected retinal organoids was significantly higher than in either day 90–infected or day 130–infected organoids (an average of 22.14 per organoid, in contrast to 8.7 for day 90 and 3.8 for day 130). Strikingly, SARS-CoV-2–infected cells appeared to localize to the inner retina in younger retinal organoids whereas they predominantly affected the outer retina in older organoids.

### Retinal ganglion cells are particularly susceptible to SARS-CoV-2 infection

To identify the retinal cells that are infected, day 74 infected organoids were co-immunostained with antibodies directed against SARS-CoV-2 N and retinal cell type–specific markers (Figure 3A-D). Interestingly, RGCs appeared to be the most commonly infected cell type, as about 40% of the N+ cells also stained positive for the RGC marker SNCG (Figure 3A and D). A clear yet smaller number of OTX2+ photoreceptors (and perhaps a few bipolar cells) and AP2A+ amacrine cells (Figure 3B and D) also stained positive for SARS-CoV-2 N, indicating the permissibility of these cells. An even smaller number of VSX2+ progenitors (and perhaps a few bipolar cells) also stained positive for N (Figure 3C and D), suggesting that progenitor cells can also be infected. As Müller glia cells are still rare at this stage of retinal organoid development, we did not attempt to stain the organoids with a Müller glia marker as well.

**Figure 3:**
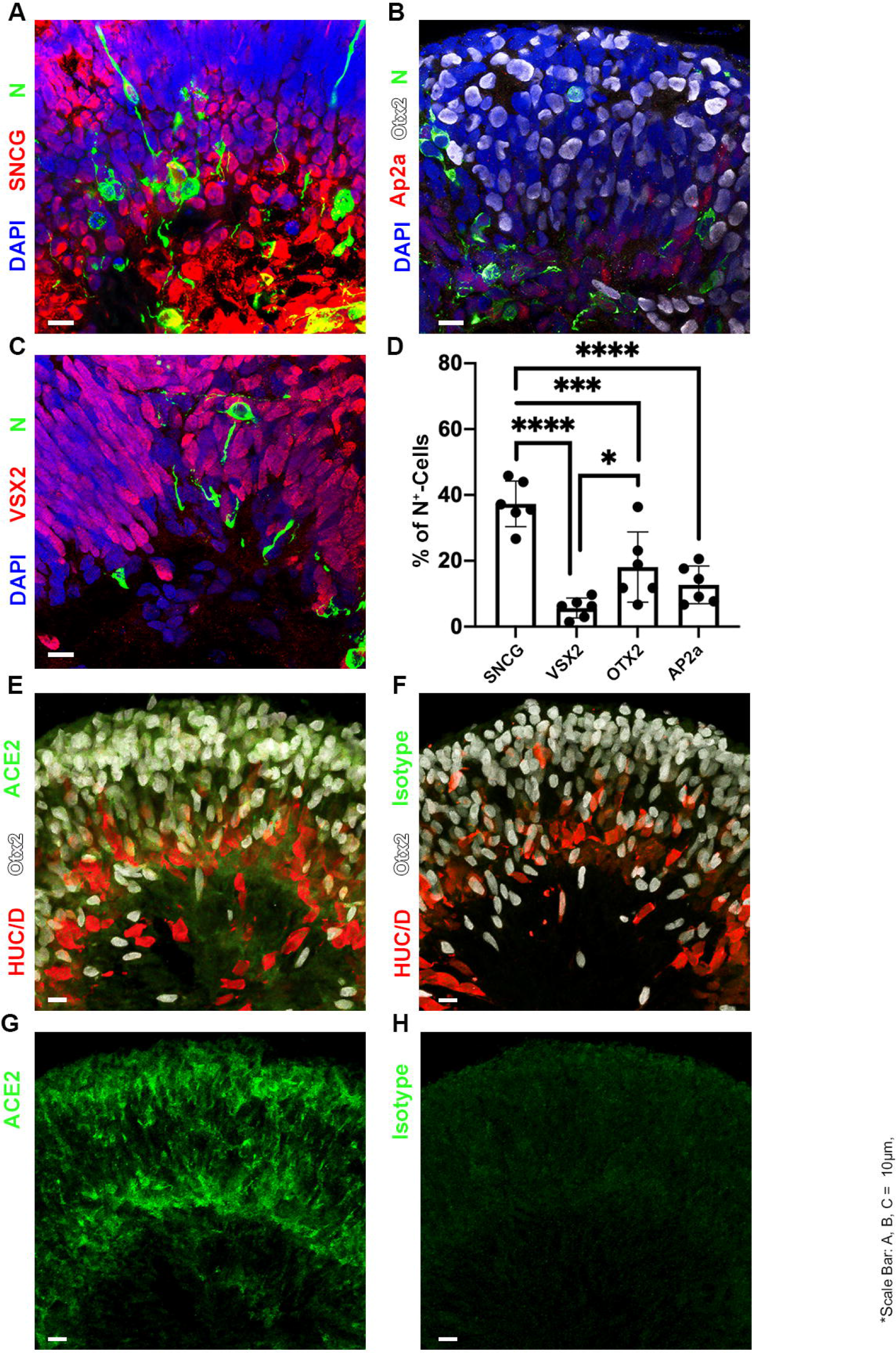
SARS-CoV-2 infects retinal ganglion cells at a higher rate than other retinal cell types. Retinal organoids infected with SARS-CoV-2 on day 74 and incubated for 96 h before fixation were immunostained with antibodies against SARS-CoV-2 nucleocapsid (N, green) and the retinal ganglion cell marker SNCG (white) **(A)**, the photoreceptor cell marker OTX2 (white) **(B)**, the amacrine and horizontal cell marker AP2α (red) **(B),** and the retinal progenitor marker VSX2 (red) **(C).** The number of cells that were co-stained with N and each of the markers was quantified, and the percentage of each marker-positive cell type out of the total number of N+ cells was compared **(D)**. N = 6 organoids taken from two different infected wells, ANOVA P-value < 0.0001, F = 21.69; ****: P-value <0.0001; ***: P-value = 0.0008; *: P-value = 0.032. Sections from day 74–control organoids were also immunostained with antibodies against ACE2 (green), together with antibodies against OTX2 (white) and HUC/D (red) **(E) and (G)**. ACE2 staining was compared to sections from the same organoids that were stained with an appropriate rabbit polyclonal antibody isotype control.

ACE2, is the receptor most commonly associated with the SARS-CoV-2 infection pathway. While ACE2 mRNA expression in the human retina is thought to be low, ACE2 protein is present in the human and rat retina (Senanayake et al., 2007; Tikellis et al., 2004). Accordingly, an immunohistochemical analysis identified an ACE2 signal concentrated in OTX2+ photoreceptors and HUC/D+ ganglion and amacrine cells in our retinal organoids (Figure 3E and G). These results are consistent with the retinal infection profile of SARS-CoV-2 shown here.

### A transcriptomic analysis of SARS-CoV-2 infected retinal organoids

To study the effect of SARS-CoV-2 infection on retinal organoids, we conducted an RNA-seq analysis to compare the transcriptome of noninfected retinal organoids to that of retinal organoids that were infected with SARS-CoV-2 on day 80 of differentiation and incubated for either 24 or 96 hours after infection. Unsurprisingly, 10 SARS-CoV-2 gene transcripts were highly enriched within the infected samples, though no difference was seen between 24 hours post-infection incubation and 96 hours post-infection incubation (Figure 4A). This is consistent with the reduction in plaque-forming units (PFU) observed in the plaque assay after 72 hours (Figure 1D).

**Figure 4:**
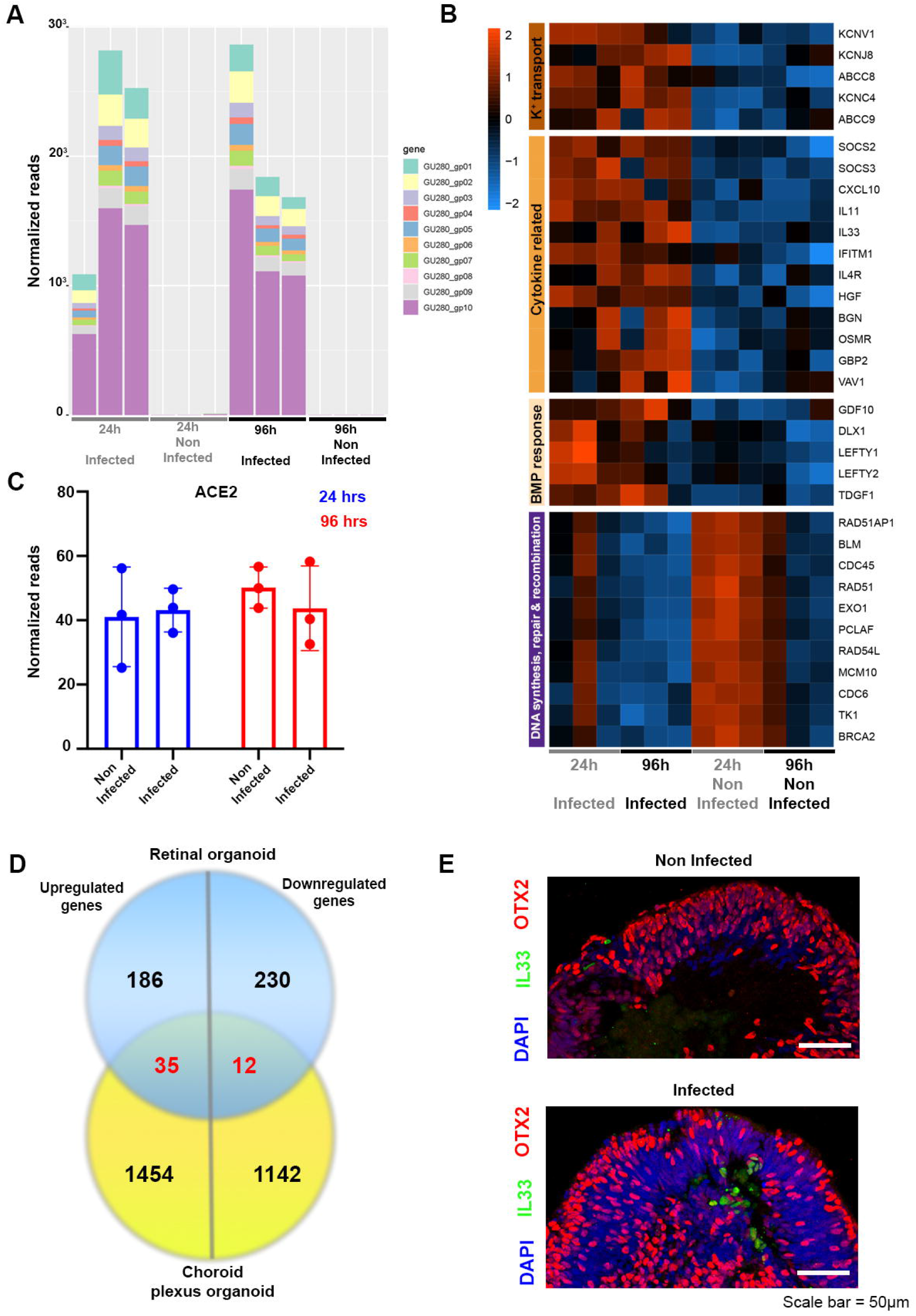
A transcriptomic analysis of SARS-CoV-2–infected retinal organoids. RNA was extracted from SARS-CoV-2 infected and control organoids on day 80 of differentiation and incubated for 24 or 96 hours following infection and from their respective controls. N=3 repeats for every treatment. Each repeat included five organoids, that were treated separately from the rest of the samples. SARS-CoV-2 transcripts were identified in large numbers in the infected samples **(A)**. Genes that were significantly differentially expressed (DE) with a fold change of at least 2 (or −2) among all of the samples together were used for gene ontology (GO) analysis using the Erichr software. A heatmap of genes related to the top enriched categories identified in the GO analysis is presented **(B).** The number of normalized reads of the ACE2 gene in noninfected and infected samples at both incubation times was also extracted from the analysis **(C)**. The DE genes identified in the retinal organoids after 24 hours were compared to the DE genes identified in SARS-CoV-2–infected choroid plexus after 24 hours. The comparison is presented as a Venn diagram **(D)**. Control and retinal organoids infected on day 74 of differentiation and incubated for 24 hours before fixation were immunostained against IL33 (green) and OTX2 (red) **(E)**.

In accordance with the immunohistochemical analysis, ACE2 transcripts were identified in all of the samples, with no change in expression due to virus infection at either incubation time (Figure 4C), suggesting that the infection of retinal cells does not result in ACE2 upregulation. This result contradicts studies suggesting ACE2 upregulation upon SARS-CoV-2 infection of other types of host cells (Butler et al., 2021).

Next, we compared the transcriptome of all of the infected samples to that of the control samples to identify genes that are differentially expressed (DE) upon infection. This analysis identified 759 DE genes (FDR adjusted P-value ≤ 0.05), of which 366 were upregulated and 393 were downregulated (Figure S1C).

To gain further information regarding the processes occurring in the retinal organoids upon infection, a gene ontology (GO) analysis of the most highly upregulated genes (FDR adjusted P-value ≤ 0.05, fold change ≥ 2) was performed using the Enrichr software (Chen et al., 2013; Kuleshov et al., 2016). The top five most enriched biological process categories in this analysis were “Potassium ion transport”, “Cellular response to cytokine stimulus”, “Cellular response to BMP stimulus”, “Metal ion transport”, and “Cytokine-mediated signaling pathway” (Figure 4B and Table 1).

The enrichment of cytokine-related genes indicates the mounting of an immune response upon SARS-CoV-2 infection. KEGG pathway analysis performed using Enrichr with the same genes found that SARS-CoV-2 infection also led to the enrichment of genes related to the JAK-STAT signaling pathway, a signaling pathway involved in inflammation and the innate antiviral interferon response (data not shown). The enrichment of potassium ion transport-related genes in the data-set may correspond with evidence suggesting that hypokalemia is a relatively common pathology of COVID-19 (Mabillard and Sayer, 2020).

Comparison of the transcriptome changes of SARS-CoV-2–infected retinal organoids to those of other neural progenitor–derived organoid systems may allow for a better understanding of the unique features of the retinal response to SARS-CoV-2 infection. To this end, we have used a data set of genes altered in choroid plexus organoids upon SARS-CoV-2 infection (Jacob et al., 2020). While the choroid plexus is not a neuronal tissue like the retina, it does develop from neural progenitors of the neural tube (Liddelow, 2015), and may share some of the response to SARS-CoV-2 infection as the retina.

Comparison of the DE genes of infected retinal organoids after 24 hours incubation to those of infected choroid plexus organoids after a similar incubation, identified 35 genes that are upregulated and 12 genes that are downregulated in both organoid systems (Figure 4D). Of the genes upregulated in both systems, 17 were highly upregulated in the retinal organoids (fold change ≥ 2). GO analysis identified enriched categories related to megakaryocyte differentiation, negative regulation of SMAD signaling and serine-threonine phosphorylation, peptidyl-tyrosine phosphorylation, and cell proliferation (Table 2). Interestingly, GO of genes that are specifically highly upregulated in the infected retinal organoids (and are not upregulated in the choroid plexus organoids) identified several enriched categories related to TGFβ and BMP signaling response (Table 2). Such a result suggests a difference in TGFβ response to SARS-CoV-2 infection between the retina and choroid plexus.

Unlike the choroid plexus organoids, retinal organoids show upregulation of the cytokine interleukin 33 (IL33), the most upregulated (Fold change = 8.11, FDR adjusted P-value = 4.27E-7) cytokine in these organoids. Accordingly, an immunohistochemical analysis identified several IL33+ cells in infected organoids, but hardly any in the controls (Figure E). Thus, SARS-CoV-2 retinal infection appears to induce IL33 expression. IL33 signaling is thought to play a role in COVID-19 pathology (Zizzo and Cohen, 2020), as well as in retinal inflammation and photoreceptor degradation following injury (Xi et al., 2016).

Another interesting inflammation-related gene found to be upregulated in SARS-CoV-2–infected retinal organoids is the inflammasome gene NLRP1 (fold change 2.2, adjusted P-value = 0.0346). NLRP1 inflammasomes are thought to be involved in RGC death in acute glaucoma (Yerramothu et al., 2018).

Interestingly, the top five categories enriched in the genes most significantly downregulated in infected retinal organoids (FDR adjusted P-value ≤ 0.05, fold change ≤ −2) are all related to DNA metabolism, repair, and recombination (Figure 4B and Table 1). This might suggest an effect of SARS-CoV-2 on retinal progenitor cells or DNA damage repair mechanisms. Other coronaviruses have been shown to interact with the cell cycle and DNA damage response in different ways (Surjit et al., 2006; Xu et al., 2011; Zhou et al., 2008).

### ACE2 blocking successfully reduces SARS-CoV-2 infection of retinal organoids

In order to assess the extent to which retinal SARS-CoV-2 infection is mediated by ACE2, retinal organoids were treated with an anti-ACE2 antibody that blocks SARS-CoV-2 infection (Hoffmann et al., 2020; Song et al., 2021). Interestingly, retinal organoids that were treated with the anti-ACE2 antibody contained a significantly lower number of SARS-CoV-2 N+ cells compared with organoids from the same batch that were treated with a normal goat IgG antibody isotype control (1.35±2.02 vs. 6.78±7.78, on average, Figure 5A, B and C). Thus, in contrast to recent suggestions that SARS-CoV-2 infection of the retina is not mediated by ACE2 (de Figueiredo et al., 2020), our results suggest that SARS-CoV-2 infection of retinal cells is dependent upon functional ACE2 receptors.

**Figure 5:**
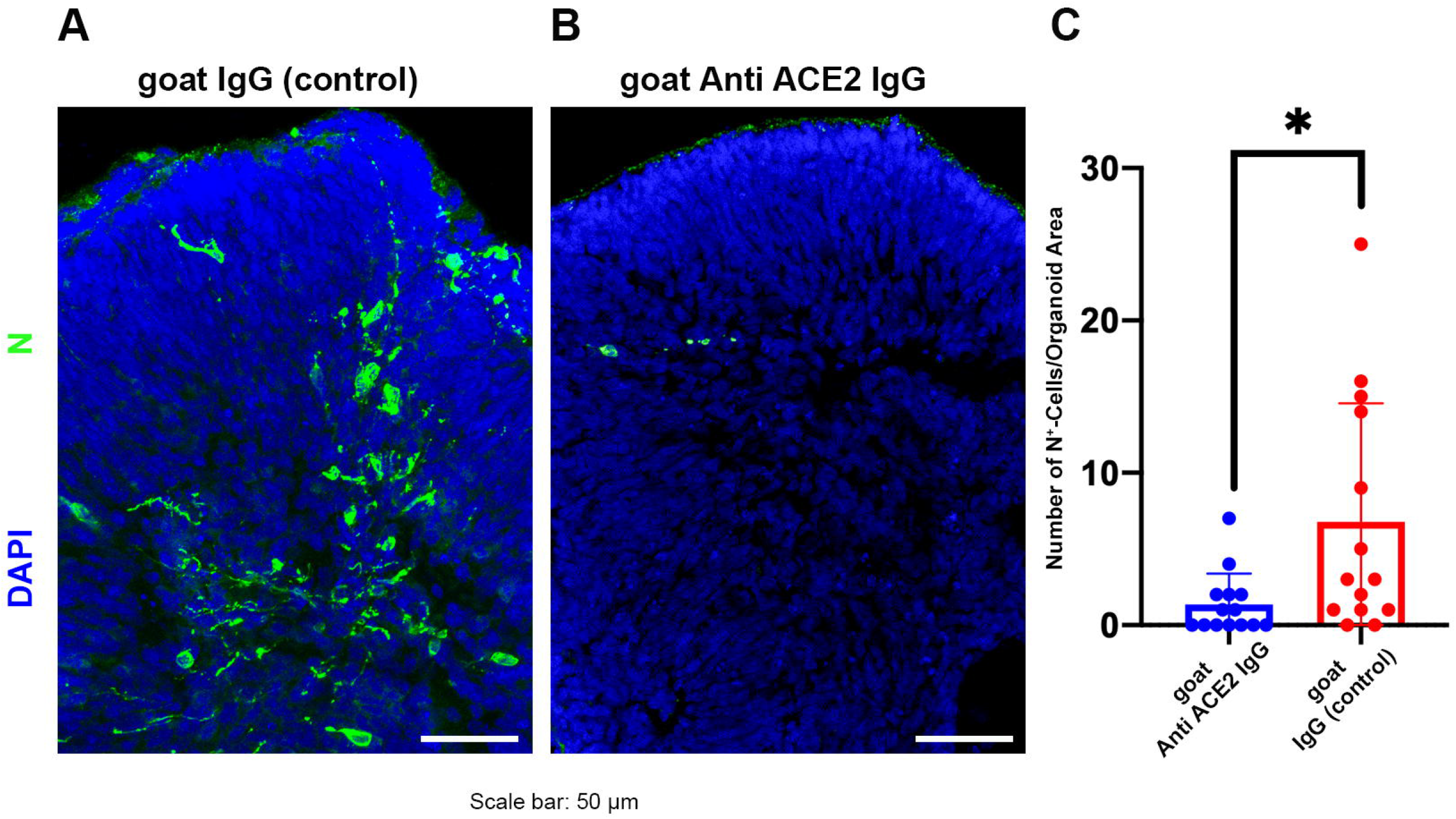
SARS-CoV-2 infection of retinal organoids is at least partially dependent upon ACE2. SARS-CoV-2 nucleocapsid (N) immunostaining (green) was used to detect SARS-CoV-2–infected cells in retinal organoids treated with either a goat IgG anti-ACE2 antibody **(A)**, or a goat IgG isotype control **(B)** before being infected with SARS-CoV-2 and incubated for 96 hours. The number of N+ cells was quantified and compared between organoids that went through the different treatments (**C)**. N = 14 organoids for each treatment, taken from three different infected wells; *: P-value (two-tailed) = 0.018.

## Discussion

### SARS-CoV-2 can actively infect retinal neurons

Retinal organoids show high similarity to the developing human retina in both morphology and transcriptome (Sridhar et al., 2020), rendering them a potent model for studying the biology of the human retina. Our data establishes retinal organoids as a potent model to study retinal involvement in COVID-19. A recently published paper showed that a recombinant lentiviral vector carrying the SARS-CoV-2 spike protein can infect retinal organoids (Ahmad Mulyadi Lai et al., 2021). Our data goes beyond that, showing that SARS-CoV-2 itself infects retinal organoids, and more importantly, we provide the first evidence that SARS-CoV-2 can actively replicate in retinal tissue.

Interestingly, younger retinal organoids appeared to be more permissive to infection than older ones. TMPRSS2 expression, which is known to be important in SARS-CoV-2 infection, was shown to be decreased in retinal organoids over time (Ahmad Mulyadi Lai et al., 2021). This might suggest that more mature retinal cells are less likely to be infected by SARS-CoV-2 because of changes in TMPRSS2 expression. Yet, the mRNA expression of TMPRSS2 in our retinal organoids was extremely low on day 80, suggesting that there is minimal involvement of TMPRSS2 in retinal SARS-CoV-2 infection.

In older retinal organoids, mature retinal cells were also infected, albeit at a lower number, indicating that mature retinal cells are still susceptible to SARS-CoV-2 infection. In fact, in the organoids, progenitor cells were less likely to be infected than differentiated neurons, emphasizing the susceptibility of retinal neurons to infection. As retinal organoids differentiate, the number of cells contained therein increases, potentially increasing the organoid density. As organoids lack vascularization, an increase in their density and size might decrease the permeability of the inner layers to SARS-CoV-2 and decrease the likelihood of viral infection of inner retinal cells.

Moreover, changes in the cell type composition might affect the infectability of the organoids. Loss of RGCs is known to occur during maturation of human retinal organoids, with nearly complete depletion of RGCs at the late stages (Sridhar et al., 2020; Zhong et al., 2014). The observed decrease in SARS-CoV-2–infected cells during retinal organoid maturation might be related to changes in the number of RGCs over time. Indeed, while our analysis identified SARS-CoV-2–infected cells from different retinal lineages, RGCs were significantly more likely to be infected.

Interestingly, many of the retinal symptoms associated with COVID-19, such as lesions in the GCL (Marinho et al., 2020) and swelling of the optic nerve (Burgos-Blasco et al., 2020), are related to RGCs. Pathology of the GCL could be the result of several issues, such as vascular dysfunction or increase in ocular pressure, and thus might be a secondary effect to other symptoms of the disease. Yet, as SARS-CoV-2 targets organoids RGCs, infection of RGCs may have such direct pathological consequences.

RGCs are the cells that generate the optic nerve and connect the retina to the rest of the CNS. Viruses, such as the herpes simplex virus type 1, can be anterogradely transferred from the retina through the optic nerve into its targets in the brain (Sun et al., 1996). Considering this, the possibility that the retina represents a potential entry route for SARS-CoV-2 into the rest of the CNS should be considered. A recent publication suggested that SARS-CoV-2 can infect many non-retinal ocular cell tissues such as the cornea, the sclera, the choroid, the limbus and the RPE, indicating the possibility of SARS-CoV-2 infection through the eye (Eriksen et al., 2021). Taken together, these results may suggest a direct route for SARS-CoV-2 infection from exposed ocular surface tissues like the cornea to the CNS through the retina.

### SARS-CoV-2 infection of retinal organoids induces inflammatory genes

The pathology of COVID-19 is highly attributed to uncontrolled hyperinflammation, referred to as a cytokine storm (Burke et al., 2020; Mehta et al., 2020). Our data suggests that the infection of retinal organoids results in the upregulation of several inflammatory genes. Among these are several genes involved in the immune response to viral infection, such as the chemokine CXCL10 (Trifilo et al., 2004) and the interferon-induced gene IFTIM1 (Brass et al., 2009), suggesting that SARS-CoV-2 infection triggers an inflammatory response.

The most significantly upregulated cytokine in infected retinal organoids was IL33. Several studies have identified a strong correlation between COVID 19 disease severity and the levels of IL33 in the blood serum of patients (Burke et al., 2020; Munitz et al., 2021). In the mature retina, IL33 is expressed mainly by Müller glia and RPE cells (Xi et al., 2016). Elevated expression and secretion of IL33 in the rodent retina were shown to be the result of different harmful stimuli such as infection with *Toxoplasma gondii*, retinal detachment, and light-induced damage (Augustine et al., 2019; Tong and Lu, 2015; Xi et al., 2016). While suggested to have a protective role in a mouse model of retinal detachment (Augustine et al., 2019), IL33 was shown to pathologically amplify the innate immune response in the rat retina following light-induced damage or RPE disruption, promoting the expression of inflammatory chemokines and cytokines from Müller glia, the accumulation of myeloid cells in the outer nuclear layer, and RGC and photoreceptor cell death (Xi et al., 2016).

Thus, our data suggests that SARS-CoV-2 infection of retinal cells results in an inflammatory response of immune system factors such as IL33 and NLRP1 which are involved in retinal degenerative diseases causing irreversible blindness. Cytokines that were up-regulated in the infected organoids, may prove to be worthy candidates for further studies regarding COVID-19 retinal pathology.

Comparison of the DE genes in retinal organoids and choroid plexus organoids identified enrichment of TGFβ response genes among genes which were induced in the retinal but not in the choroid plexus organoids, suggesting that SARS-CoV-2 infection may induce differential TGFβ response in different systems.

The eye is an immune-privileged organ, as different mechanisms reduce its inflammatory responses in order to prevent vision impairment. TGFβ signaling has an important role in maintaining the immune-privileged status of the eye (Masli and Vega, 2011; Zhou and Caspi, 2010). For example, both RPE cells and RGCs suppress the activation of T cells via the secretion of TGFβ molecules (Edo et al., 2020; Sugita et al., 2011). Unlike most of the CNS, the choroid plexus is not an immune-privileged structure (Galea et al., 2007). Thus, differences in TGFβ response of retinal and choroid plexus organoids to SARS-CoV-2 infection may be related to the immunosuppressive role of TGFβ.

### SARS-CoV-2 retinal infection is dependent on ACE2

The relatively low expression of ACE2 in the retina has led researchers to speculate that SARS-CoV-2 retinal infection is dependent upon alternate pathways (de Figueiredo et al., 2020). Yet, ACE2 protein was identified in human retinas and retinal organoids (Ahmad Mulyadi Lai et al., 2021; Senanayake et al., 2007). Our data indicates that the infection of retinal cells can be blocked by using anti-ACE2 antibodies. Thus, the infection of retinal cells most likely is primarily dependent upon functional ACE2 receptors. The possibility of blocking retinal infection by using ACE2 antibodies may also be helpful for the development of drugs in the treatment of SARS-CoV-2 retinal infection.

To conclude, our data indicates that SARS-CoV-2 can actively infect retinal cells in an ACE2-dependent manner. Further studies should consider the possibility that neuro-retinal infection leads to retinal symptoms in patients with COVID-19 and perhaps examine potential treatment options. SARS-CoV-2–dependent photoreceptor and/or RGC degeneration can cause permanent visual impairment or even blindness. While data suggesting vision impairment in patients with COVID-19 is scarce, it should be noted that visual impairment and subsequent blindness from retinal degenerative diseases may become evident only after a long course of progression. Long-Covid- or Post-Covid-Syndrome–related vision impairment or even blindness may occur at a much later time point after an acute SARS-CoV-2 infection. The induction of inflammatory genes that are related to retinal degeneration in the organoids should prompt further investigation of the association between SARS-CoV-2 infection and retinal degenerative diseases.

While our study indicates that SARS-CoCV-2 can infect and replicate in retinal cells in organoids, our experimental system has some limitations. Retinal organoids contain some RPE tissue together with neuro-retinal tissue but are devoid of other important tissues such as the cornea and the retinal vasculature present in the physiological environment (Achberger et al., 2019). This limits the ability to use the retinal organoid model to study the entry and route of SARS-CoV-2 infection in the retina and the interaction between the retina and other tissues during infection. Moreover, retinal organoids lack specialized immune cells such as microglia, which have an important role in retinal inflammation (Rashid et al., 2019), and thus cannot be used to fully characterize the retinal inflammatory response to SARS-CoV-2. Retinal organoids also represent the embryonic retina rather than a mature retina, and thus some age-related differences might affect retinal SARS-CoV-2 infection and pathology. Yet, the fact that SARS-CoV-2 also does infect relatively mature neurons and cause the upregulation of inflammatory genes in our organoids, combined with the growing evidence of retinal involvement in patients with COVID-19, indicates the relevance of the data produced from our retinal organoids and the need for further investigations into COVID-19–related retinal pathologies.

## METHODS

### Organoid production

H9 human embryonic stem cells (HESCs) and Gibco episomal iPSCs (line A18945) were grown in feeder-free conditions in Matrigel (Corning)-coated plates in StemFlex medium (Thermo Fisher). To produce organoids, we have combined different widely used retinal organoid generation protocols (Capowski et al., 2019; Kuwahara et al., 2015; Zhong et al., 2014). In short, HESCs or IPSCs were grown until they reached ~80% confluence. At that stage, cell colonies were detached from the plate using 0.5 mM EDTA and placed in a low attachment 6-well plate (MERK) well in 1 mL of mTeSR1 medium (Stem cell technologies) containing 10 μM of Blebbistatin (Sigma-Aldrich) per well. Over the next three days, the medium was replaced gradually to neural induction medium (NIM) comprising DMEM/F12 (Thermo Fisher) 1:1, 1% N2 (Thermo Fisher), 1 x MEM nonessential amino acids (Thermo Fisher), 1 x Glutamax (Thermo Fisher), and 2 μg/mL of heparin (Sigma-Aldrich). On day 6 of differentiation, NIM was replaced and supplemented with 1.5 nM of BMP4 (R&D Systems). Half of the media was replaced on days 9, 12, and 15 of differentiation with fresh NIM to dilute the BMP4 concentration. On day 16, NIM was replaced with retinal differentiation medium (RDM) comprising DMEM:F12 3:1, 2% B27 (Thermo Fisher), 1x MEM nonessential amino acids, 1x Penicillin-Streptomycin (Thermo Fisher), and 1x Glutamax. From this stage onward, the RDM medium was changed twice a week. On day 30 of differentiation, RDM was supplemented with 10% FBS and 100 μM of taurine (Sigma-Aldrich). Between days 40 and day 100 of differentiation, RDM was also supplemented with 1 μM of all-trans-retinoic acid (Sigma-Aldrich).

### Virus production and organoid infection

SARS-CoV-2 was isolated from a patient in South Tyrol (hCoV19/Germany/FI1103201/2020) and identified as type “Ischgl”. The virus sequence is available in the GSAID data bank under EPI-ISL_463008)

All SARS-CoV-2–containing experiments were conducted under biosafety level (BSL) 3 conditions. The virus was propagated on VeroE6-TMPRSS2 cells using DMEM (Sigma) supplemented with 2% (v/v) FBS (Capricon Scientific), 1% (v/v) Penicillin/Streptomycin (Sigma), 1% (v/v) sodium pyruvate solution (Sigma), 1% (v/v) NEAA (Sigma), and 1% (v/v) HEPES solution (Sigma) using an MOI of 0.01. Two days post-infection, the virus titer was determined.

SARS-CoV-2 was diluted to the desired amounts in RDM supplemented with 100 μM of taurine (Sigma-Aldrich) without FBS. The organoids were washed twice with PBS and incubated for 1 h at 37°C with the virus dilutions. Afterward, organoids were washed once with PBS and incubated in RDM supplemented with 100 μM of taurine for specific time points.

### Plaque assay

VeroE6 cells were grown to a confluent monolayer in 6-well plates. 10-fold dilution series of viral solutions were prepared in PBS (Sigma) containing 1% (v/v) Penicillin/Streptomycin, 0.6% (v/v) BSA (35%) (Sigma), 0.01% (w/v) CaCl2 (1%), and 0.01% (w/v) MgCl2 (1%). The VeroE6 cells were incubated with the dilution series for 1h at 37°C. Afterwards the inoculum was replaced by plaque medium (63% (v/v) 2x MEM [20% (v/v) 10x MEM (Gibco), 3.2% (v/v) NaHCO3 (Lonza), 2% (v/v) HEPES (1M; pH 7.2) (Sigma), 1.2% (v/v) BSA (35%), 1% (v/v) 100x Penicillin/Streptomycin/L-Glutamine solution (10.000 U/ml Pen.; 10.000 μg/ml Strep., 29.2 mg/ml L-Glutamine) (Gibco)], 2% (v/v) FBS, and 35% (v/v) Agar (2%) (Oxoid)). After 48 h to 96 h, plaques were counted.

### Immunohistochemistry

Organoids were fixed in 4% PFA for 30 minutes (organoids used for ACE2 immunohistochemistry were fixed for 2 hours), washed three times in PBS, and incubated overnight at 4°C in 30% sucrose in PBS. The organoids were then cryopreserved in O.C.T (Tissue-Tek) and sectioned to a thickness of 20 μm. The sections were blocked and permeabilized with 10% normal donkey serum (Abcam) and 0.3% Triton X in PBS for 1 hour at room temperature (RT), and then incubated with the primary antibodies diluted in blocking medium overnight at 4°C. The primary antibodies used for the study can be found in table S1. Sections were further washed and incubated with a species-specific secondary fluorescent antibody, together with DAPI, for 1 hour in the dark at RT and then mounted. Imaging was done using an LSM 780 scanning confocal microscope (Zeiss). All images presented here are maximal projection images. Cell counting was performed manually on random sections from different organoids, with a similarly sized area.

### RNA extraction and cDNA library preparation and RNA sequencing

RNA from mock-treated and infected organoids was extracted with TRIzol^®^ reagent according to the manufacturer’s protocol with minor modifications. Briefly, washed retina organoids were resuspended in 700 μL of TRIzol reagent. The tissue was disrupted mechanically by vortexing for 30 sec and pipetting 20 times. 140 μL of chloroform was added and samples were mixed vigorously for 15 sec. Following centrifugation at 4°C, the aqueous phase was carefully separated and RNA was further precipitated by 350 μL of 2-propanol. RNA pellets were washed twice with 75% ethanol and resuspended in DEPC-treated water for further library preparation.

600 ng of total RNA was used for RNA isolation, fragmentation, and cDNA synthesis using the NEBNExt^®^ Poly(A) mRNA magnetic isolation module (E7490). cDNA libraries were amplified and index labeled using the NEBNext^®^ Ultra TM II RNA Library Prep kit for Illumina^®^ (E7770, E7775). DNA library quality and concentration were analyzed on an Agilent Bioanalyzer DNA Chip. cDNA samples were loaded on a NextSeq 2000 system (Illumina) at a concentration of 800 pMol, and the sequencing was performed with single-end 100-bp reads.

### Bioinformatic analysis

The sequencing data was first demultiplexed using the Illumina software bcl2fastq v2.20.0. The quality of the resulting FASTQ files was then evaluated with the program FastQC v0.11.8 (https://www.bioinformatics.babraham.ac.uk/projects/fastqc/).

To perform the alignment, a customized combination of the human genome and the SARS-CoV-2 viral genome was created. The reference sequence and gene annotation for SARS-CoV-2 were obtained from the NCBI repository, from the entry with accession number NC_045512.2 “Severe acute respiratory syndrome coronavirus 2 isolate Wuhan-Hu-1, complete genome”. The human genome sequence and gene annotation were also obtained from NCBI, using the assembly version GRCh38.p13. Both viral and human genomes were concatenated using a customized bash script to create the alignment reference sequence and annotation files. Finally, the alignment was performed using STAR v2.7.7a (Dobin et al., 2013) with default parameters and the extra instruction “–-quantMode GeneCounts” to create additional gene count tables.

The following secondary analysis was performed in the statistical environment R v4.0.3 (https://www.r-project.org/). After importing the gene count tables, only genes that had a minimum of 5 reads in at least 3 samples were kept as a quality filter. Then, a dispersion trend among the samples was assessed with a principal component analysis (PCA) applied over the normalized read counts, transformed through the regularized log transformation (Figure S1B). Differential gene expression was assessed based on a Negative Binomial distribution through the package DEseq2 v1.30.1 (Love et al., 2014). Genes were considered as differentially expressed (DE) if they presented an absolute value of fold change of >1 and a Benjamini-Hochberg FDR of ≤ 5%. A very small number of SARS-CoV-2 transcripts was identified in one of the three 24-hour control samples, yet considering the vast difference in scale between the levels of SARS-CoV-2 transcripts in this sample and the levels found in the infected samples (Figure 4A, fold change > 200), these transcripts most likely do not represent a SARS-CoV-2 infection in this sample. In addition, this sample clustered together with the other control samples in a PCA analysis (Figure S1B). Thus, we decided to include this sample in the analysis.

Heatmap representation of the expression of selected genes was created with the package pheatmap v1.0.12 using the values obtained after applying the regularized log transformation over the raw counts.

GO and KEGG pathway analyses were performed by separately collecting all upregulated genes with a fold change of ≥2 and all downregulated genes with a fold change of ≤ −2 and uploading the gene lists to the Enrichr software (Chen et al., 2013; Kuleshov et al., 2016).

### Real-time PCR

cDNA synthesis was performed using the High-Capacity cDNA Reverse Transcription Kit (Applied Biosystems). Real-time PCR was performed using iTaq SYBRGreen Supermix with ROX (Bio-Rad) and a QuantStudio3 machine (applied biosystems). Gene expression was normalized according to the housekeeping gene GAPDH and calculated using the delta CT method. Primers used in this research can be found in table S2.

### ACE2 blocking

Retinal organoids on day 90 of differentiation were incubated at RT for 1 hour in PBS with either a goat anti-ACE2 antibody (AF933, R&D Systems) or a normal goat IgG control (AB-108-C, R&D Systems), at a concentration of 100 μg/mL before being infected with SARS-CoV-2.

### Statistical analysis

Statistical analysis was done using GraphPad Prism 9. Analysis of experiments with multiple groups was done by a one-way ANOVA with Tukey’s multiple comparison test. Statistical analysis of experiments with only two groups was done using an unpaired, two-tailed Student’s t-test. No statistical tests were used to predetermine the sample size.

## Data availability

Raw and processed RNA-seq data was uploaded to GEO.

## Acknowledgements

This research was supported by the Max Planck Society’s *White Paper-Project: Brain Organoids: Alternatives to Animal Testing in Neuroscience*. Further financial support was received from the German Research Foundation (DFG), grant SFB1009 B13, and the German Ministry of Education and Research (BMBF), project OrganoStrat (01KX2021), SFB944 (Z-Project) and the IZKF, grant Bru2/015/19. We thank Ingrid Gelker and Manuela Haustein for their technical help. We also thank the team of the Core Facility Genomik in the medical faculty in Münster for RNA-sequencing and Areti Malapetsas for editing the manuscript.

## Author contributions

YML, AS, SL, TR and HRS conceived the study.

YML and TR wrote the manuscript and prepared the figures.

YML, AS, AMZ, LB and OEP performed experiments.

AL analyzed data

All authors contributed to the editing of the manuscript

---

**Figure S1: Supplementary information to figures 1 and 4**

A retinal organoid derived from H9 embryonic stem cells stained with an anti–SARS-CoV-2 nucleocapsid (N) antibody (green). Scale bar = 50 μM. **(A)**. PCA analysis of the different infected on noninfected samples was performed using the rlog algorithm **(B)**. MA plot representing the mean expression levels and the fold change of the different genes between the infected samples and the noninfected samples. Significantly differentially expressed genes (adjusted P-value ≤ 0.05) are colored red **(C)**. Venn diagram comparing the differentially expressed genes found in retinal organoids 96 hours post-infection and the differentially expressed genes found in choroid plexus organoids after 72 hours. Only 7 genes are mutually upregulated and 4 genes are mutually downregulated **(D)**.

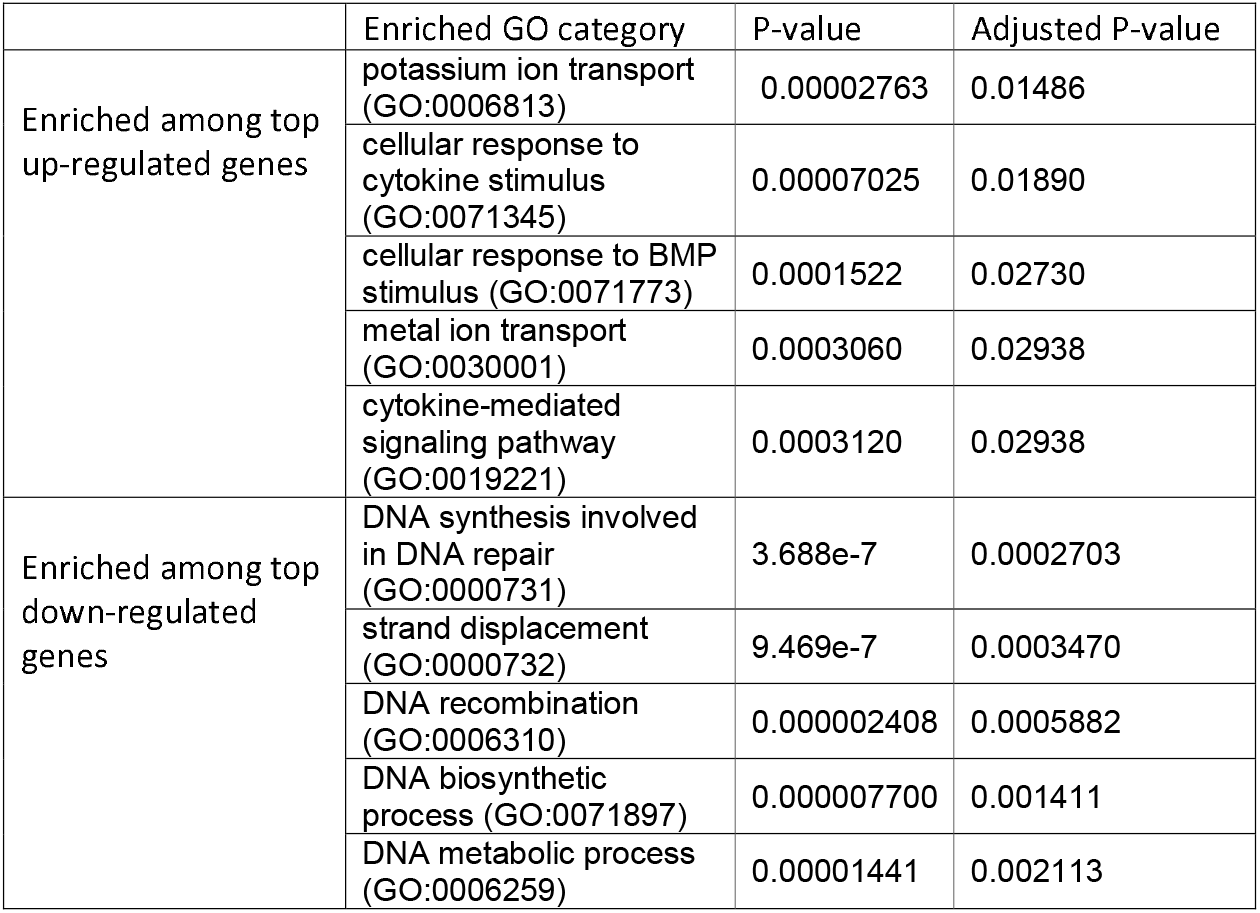

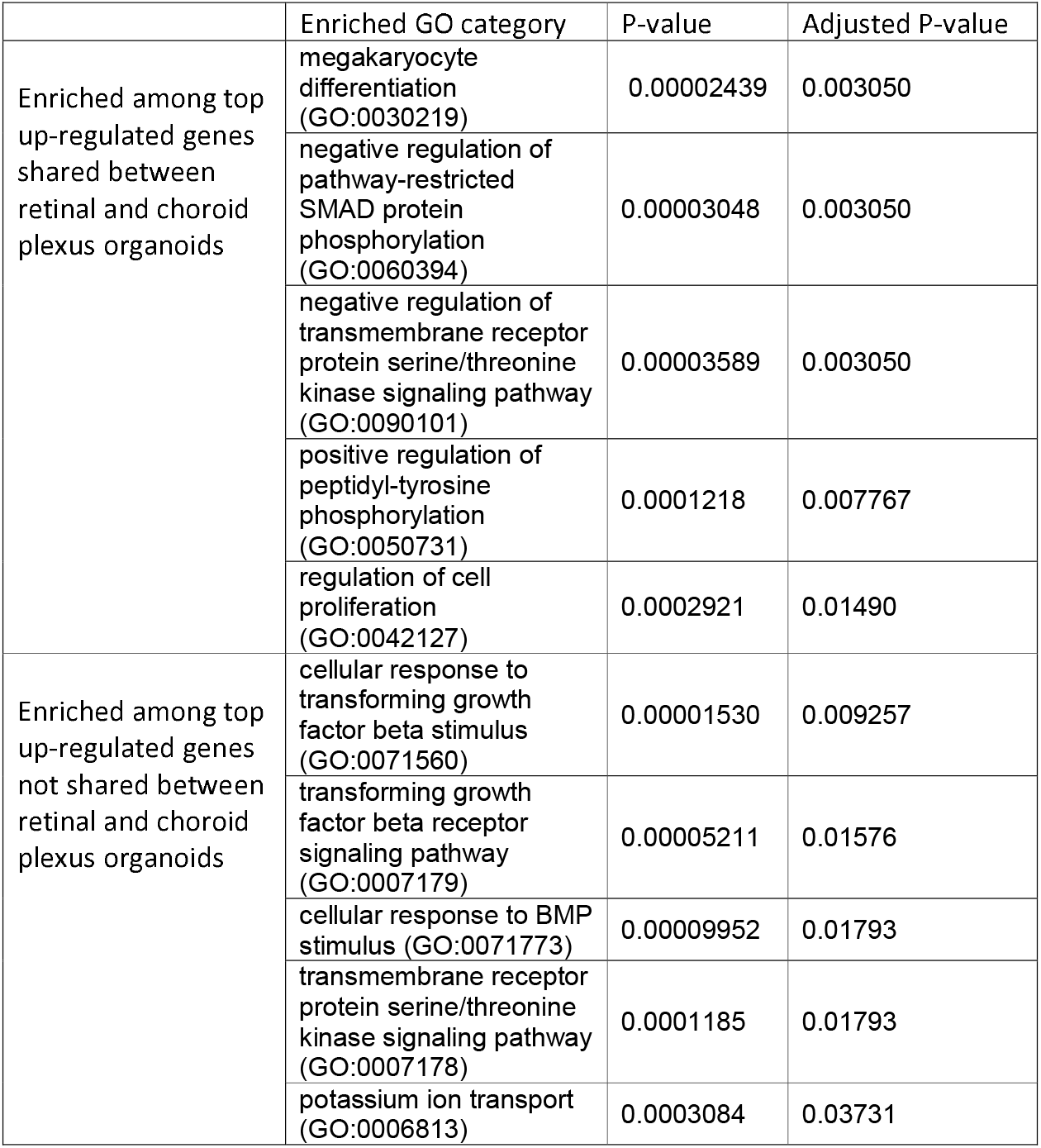

