## Supplementary material for "SARS-CoV-2 infects and replicates in photoreceptor and retinal ganglion cells of human retinal organoids": Table S1 and S2

| **Antibody** | **Source** | **Cat. num** | **Dilution** |
| --- | --- | --- | --- |
| ACE 2 | Abcam | ab1348 | 1:500 |
| AP2A | Abcam | ab220065 | 1:200 |
| CHX10 (VSX2) | Milipore | ab9014 | 1:500 |
| CRALBP | Abcam | Ab15051 | 1:500 |
| HUC/D | Invitrogen | A21271 | 1:100 |
| Rabbit IgG polyclonal isotype control | Abcam | Ab37415 | 1:500 |
| IL33 | Abcam | Ab118503 | 1:200 |
| L/M Opsin | Milipore | AB5405 | 1:500 |
| OTX2 | R&D systems | AF1979 | 1:500 |
| PAX6 | Biolegend | 901301 | 1:500 |
| RHODOPSIN | Millipore | MAB5316 | 1:500 |
| SARS-CoV/SARS-CoV-2 nucleocapsid | Sino biological | 40143-MM05 | 1:200 |
| SNCG | Abcam | Ab55424 | 1:500 |

**Table S1 –** primary antibodies used for the study

| Target | Forward | Reverse |
| --- | --- | --- |
| GAPDH | TGATGACATCAAGAAGGTGGTG | ACCCTGTTGCTGTAGCCAAT |
| SARS-CoV-2 | CGCATACAGTCTTRCAGGCT | GTGTGATGTTGAWATGACAT |

**Table S2 –** Primers used for the study
