## Supplementary figures and images for "SARS-CoV-2 infects and replicates in photoreceptor and retinal ganglion cells of human retinal organoids"

### Figure S1

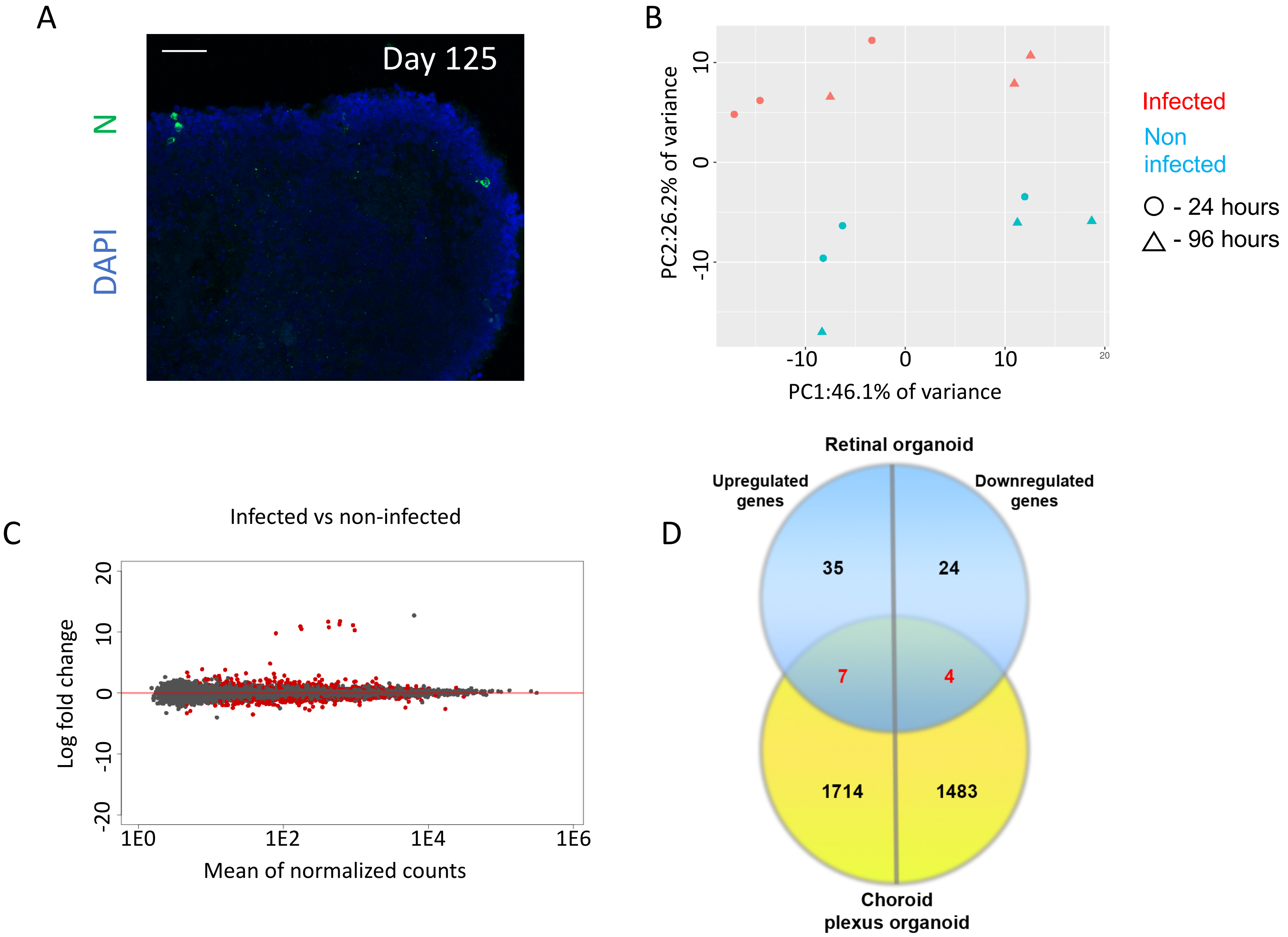
